# A multi-level account of hippocampal function from behaviour to neurons

**DOI:** 10.1101/2022.06.09.495367

**Authors:** Robert M. Mok, Bradley C. Love

## Abstract

A complete neuroscience requires multi-level theories that address phenomena ranging from higher-level cognitive behaviors to activities within a cell. A levels-of-mechanism approach that decomposes a higher-level model of cognition and behavior into component mechanisms provides a coherent and richer understanding of the system than any level alone. Toward this end, we decomposed a cognitive model into neuron-like units using a *neural flocking* approach that parallels recurrent hippocampal activity. Neural flocking coordinates units that collectively form higher-level mental constructs. The decomposed model suggested how brain-scale neural populations coordinate to form assemblies encoding concept and spatial representations, and why so many neurons are needed for robust performance at the cognitive level. This multi-level explanation provides a way to understand how cognition and symbol-like representations are supported by coordinated neural populations (assemblies) formed through learning.

## Introduction

Neuroscience is a multi-level enterprise. Its target of explanation ranges from behavioral to molecular phenomena. Satisfying and complete explanations of the mind and brain will necessarily be multi-level (*1–3*). In multi-level componential (or constitutive) explanations, each component at a higher level can be decomposed into its own lower-level mechanism (*1*). For example, the circulatory system’s capacity to deliver oxygen and energy to the body can be decomposed into lower-level mechanisms including the heart’s blood pumping and kidney’s blood filtering mechanism, which together supports the function of the circulatory system. These mechanisms themselves can be decomposed into their components, such as the muscle contractions of the heart or filtering units of the kidney, which in turn can be further decomposed as desired (*1*).

In neuroscience, the highest-level mechanism could be a cognitive model that captures behaviour. The components of this cognitive model could be related to neural measures and further decomposed. What is a component at a higher-level is a mechanism at a lower-level that can itself be decomposed into components to account for additional findings and make new predictions. The power of this approach is that mechanisms at different levels are not unrelated efforts aiming to explain different types of data. Instead, multi-level explanations can be integrated and offer a more complete understanding of the domain in which one can “zoom in or out” to the level of granularity desired for the current question.

Constructing theories and testing their predictions at multiple levels provides a more comprehensive account of the system of interest. Although neuroscience is guilty of a bias toward lower-level explanations (*4, 5*), higher-level mechanisms are crucial in multi-level explanation because they offer explanatory concepts not available at lower levels (*3*). For example, the heart’s contractions make little sense without considering the function of the circulatory system, and the hippocampal synaptic weights and activity patterns make little sense without notions of memory and learning. In neuroscience, it is common to construct a specific theory or model to fit the specific data at hand, which unfortunately leads to a disconnected patch-work of theories for individual phenomena. Multi-level theories can weave this patch-work together into a coherent and complete account in which each level is situated within mechanisms that lie above and below. As one descends levels, additional phenomena can be addressed, whereas as one ascends the function of the mechanism within the overall system becomes clearer.

Neuroscience has very few multi-level theories of this variety that can bridge between cognitive constructs and neuronal activity – multi-level explanations from behavior to neurons. For example, how does the brain implement a symbol? Specifically, how do brain systems coordinate neural populations to form symbol-like representations, such as highly selective neural assemblies (*6–8*) that encode concepts (*9*) or specific spatial locations (*10*)? One suggestion is that the hippocampal formation represents physical space (*11–13*) as well as abstract spaces for encoding concepts and abstract variables (*14–16*), constructing cognitive maps (*17*) for mental navigation in these spaces. However, it is unclear how populations of similarly-tuned neurons in the hippocampus acquire their highly-selective tuning properties to concepts or spatial locations. While tantalising, identifying a cell tuned to particular concept, such as Jennifer Aniston (*9*), does not specify the supporting mechanisms leaving such findings as curiosities that invite explanation.

Attempts have been made to offer multi-level theories in neuroscience but critical explanatory gaps remain. In our own work, we have developed a cognitive model, SUSTAIN, of how people learn category from examples. SUSTAIN addresses a number of behavioural findings (*18, 19*) and aspects of the model have been related to the brain (*19–22*). The hippocampus was related to SUSTAIN’s *clustering* mechanism, which bundles together relevant information in memory during learning, and this account was verified by a number of brain imaging studies (*19–21*). The goal-directed attentional mechanism in SUSTAIN was linked to the ventral medial prefrontal cortex and this account was verified by brain imaging (*22*) and animal lesion studies (*23*). These same mechanisms provide a good account of place and grid cell activity in spatial tasks (*16*).

Although wildly successful in addressing both behaviour and accompanying brain activity, this line of work, like almost all work in model-based neuroscience, is limited to (1) proposing correspondences between model components and brain regions and (2) evaluating these correspondences in terms of model-brain correlates. But how do we move beyond mere neural correlates to a lower-level mechanistic explanation that unpacks the higher-level theory? Cognitive models, such as SUSTAIN, come with abstract constructs such as clusters, and it is left entirely open how they could be decomposed into the neural populations that give rise to behavior. It is insufficient to state that each cognitive construct (e.g., a cluster) is instantiated by a number of neurons, just as it is unsatisfying to state that Jennifer Aniston is somehow represented by multiple neurons, the spreadsheet on a computer relies on a number of transistors, and so forth. How do neurons coordinate to give rise to the higher-level construct? This is the key question that needs to be addressed to move beyond mere neural correlates toward a mechanistic understanding of how the brain implements cognition.

We aim to address this explanatory gap by decomposing aspects of a cognitive model, SUSTAIN, into a mechanism consisting of neuron-like units. Critically, the aggregate action of these neuron-like units give rise to virtual structures akin to the higher-level cognitive constructs in SUSTAIN (i.e., clusters) while retaining SUSTAIN’s account of behavior (Fig. 1). By taking a key component of a cognitive model that addresses behaviour and decomposing it to the level of a neuron, we offer an account of how concept cells and spatially-tuned cell assemblies can arise in the brain.

**Fig. 1.**
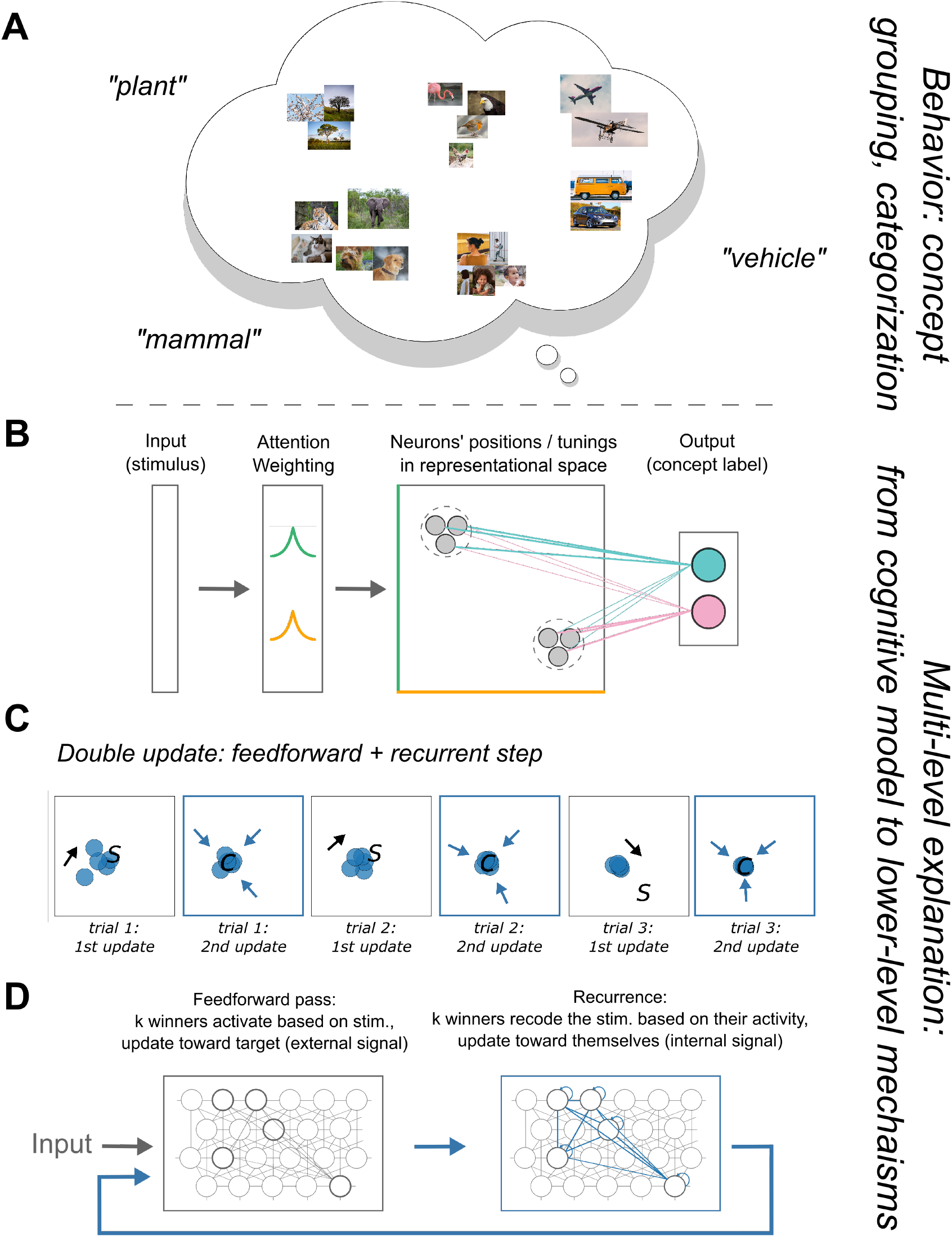
Multi-level explanation of concept learning in the brain: decomposition of a cognitive model into neural flocks. A) Concept learning and representation: the task. B) Model illustration. After the stimulus is encoded, attention is applied and neuron-like units activate according to their similarity to the input. These activations are transmitted through learned association weights to generate an output (e.g., category decision). Grey circles represent neuron-like units, dotted circles highlight units in the same flock or virtual cluster. C) Illustration of the flocking learning rule. Left to right: *k* winners (blue) move toward the stimulus (“S”), followed by a recurrent update, where units move towards their centroid (“C”). Over time, *k* neuron-like units become similarly tuned, forming a neural flock. D) Neural flocking parallels two forms of hippocampal recurrence. The *k* most activated neuron-like units (dark grey circles) not only adjust their receptive fields to move toward the stimulus but also move toward each other, implementing flocking. In effect, recurrence re-codes the stimulus in terms of the winners’ response (blue box, blue circles; lines represent connections). An alternative and complementary mechanism is big-loop recurrence in which the winners’ response is recirculated as input, akin to hippocampal output being fed back into the hippocampus via entorhinal cortex (blue arrow looping back to input).

One of the main challenges is how to bridge from abstract constructs such as clusters to neurons, whilst retaining the high-level behavior of the model. How do single neurons track a concept? How does the brain coordinate the activity of many neurons during learning, as there are thousands of neurons, but only a few clusters are required to represent a concept at the cognitive level? In other words, how does a select population of neurons learn and become tuned to similar features in the world such as in concept cells and place cells, rather than independently develop their own tuning? To implement a higher-level construct, such as a cluster or symbol, neurons must somehow solve this *coordination problem*.

Inspired by algorithms that capture flocking behavior in birds and fish (*24*), we propose the hippocampus may exhibit *neural flocking*, where coordinated activity – virtual clusters or flocks – arise from local rules (Fig. 1). This coordination could be achieved by recurrent processing in the hippocampus (e.g., (*25*)), which we formalise in learning rules in which neuron-like units that are highly active in response to a stimulus (e.g., a photo of Jennifer Aniston) both adjust their tuning toward that stimulus and to each other. The latter learning update is the key to flocking as members of the neural flock coordinate by moving closer to each other. That simple addition to standard learning rules is sufficient to solve the coordination problem and give rise to virtual clusters as well as hippocampal place or concept cell (*9*) assemblies (*7*).

Gazing at the neuron-like units forming the model, one will not see clusters just as one will not see clusters nor symbols by peering into the grey goo of the brain. Nevertheless, the coordinated activity of these neuron-like units can be described as supporting these higher-level constructs that behave in aggregate like the higher-level cognitive model, SUSTAIN. In the model specification and simulations that follow, we aim to close the aforementioned explanatory gap and make the case for multi-level explanation in neuroscience. Additionally, by decomposing the higher-level model we can consider how the brain benefits in terms of fault and noise tolerance by implementing clusters in a large neuronal flock. For instance, the mental representation of a concept is preserved when one neuron, or even a set of neurons in the neural assembly, dies, as well as when there is significant synaptic turnover over relatively short time scales (typical in the hippocampus; (*26*)). Finally, we consider how the model can be extended and further decomposed to account for differences in processing across anterior-posterior hippocampus axis.

## Multi-neuron clustering model

Our multi-neuron clustering model, SUSTAIN-d, is a decomposition of the SUSTAIN model of category learning. Whereas prototype models always form one unit in memory for each category and exemplar models store one unit for each episode, SUSTAIN moves between these two extremes depending on the nature of the learning problem. SUSTAIN assumes the environment is regular and clusters similar experiences together in memory until there is a surprising prediction error, such as encountering a bat and wrongly predicting that it is a bird based on its similarity to an existing bird cluster. When such an error occurs, a new cluster is recruited. Notable episodes (e.g., encountering a bat for the first time) can transition to concepts over time (e.g., other similar bats are stored in the same cluster). The hippocampus is critical in supporting this form of learning (*27*). SUSTAIN has other elements that are not the focus of this contribution, such as an attentional mechanism that determines which aspects of stimuli are most relevant to categorisation.

Here, we decompose SUSTAIN into more neuron-like units while retaining its overall function. We will refer to this model as SUSTAIN-d for SUSTAIN decomposed (see Methods for formal model description). Rather than recruit cognitive units like clusters, SUSTAIN-d, like the hippocampus, has an existing pool of neuron-like computing that enter each task with some tuning (i.e., preferentially activated for particular stimuli). Unlike SUSTAIN, which will recruit a handful of clusters for a learning problem, SUSTAIN-d can consist of an arbitrarily large number of computing units (see Results for brain-scale simulations where the number of computing units is equal to the number of neurons in the hippocampus).

Despite these striking differences, SUSTAIN-d’s thousands of neuron-like computing units show the same aggregate behaviour as SUSTAIN. This is accomplished by solving the afore-mentioned coordination problem by what we refer to as neural flocking (Fig. 1c). The key to neural flocking is that units that are highly activated by a stimulus both adjust their tunings toward the stimulus and each other. This double update leads to virtual clusters forming that can consist of thousands of neuron-like computing units. In general, the number of clusters SUSTAIN recruits will match the number of neural flocks that arise in SUSTAIN-d, which leads to the models providing equivalent behavioural accounts (Fig. 1A). Whereas SUSTAIN associates clusters with a category or response, SUSTAIN-d’s individual neuron-like units form connection weights (Fig. 1B). In summary, SUSTAIN-d is formulated absent of cognitive constructs like clusters, but nevertheless its neuron-like units in aggregate manifest the same principles and can account for the same behaviours, which provides an account of how cognitive constructs can be implemented in the brain.

## Formation of concept and spatial representations by neural flocking

SUSTAIN-d’s neural flocking mechanism can explain how concept cells (Fig. 2A-B) and spatially-tuned place and grid cells arise (Fig. 2C-D). Whereas our previous work (*16, 18*) relied on higher-level mental constructs (i.e., clusters) to account for such phenomena, here we show how a neural population can coordinate to virtually form such structures via the flocking learning rule. In the spatial domain, we simulated an agent (e.g., rodent) exploring a square environment (free foraging), which leads to virtual clusters akin to place cells distributed in a grid-like pattern (Fig. 2C), which in turn leads to cells that monitor these units’ activity displaying a grid-like firing pattern (Fig. 2D). In the conceptual domain, where the representation space is not as uniformly sampled, virtual clusters are clumpier (Fig. 2B) and monitoring units will show no grid response. These results in the conceptual and spatial domain hold across a wide-range of parameters. The flocks that arise from the interaction of numerous neuron-like units are a higher-level representation in that the number of underlying neuron-like units involved can vary over orders of magnitude with the aggregate behavior at the “flock level” remaining constant. This robustness allows SUSTAIN-d to decompose cognitive constructs to neuron-like units at same scale as the brain as demonstrated by simulations where the number of units is equal to the number of neurons in the corresponding brain regions.

**Fig. 2.**
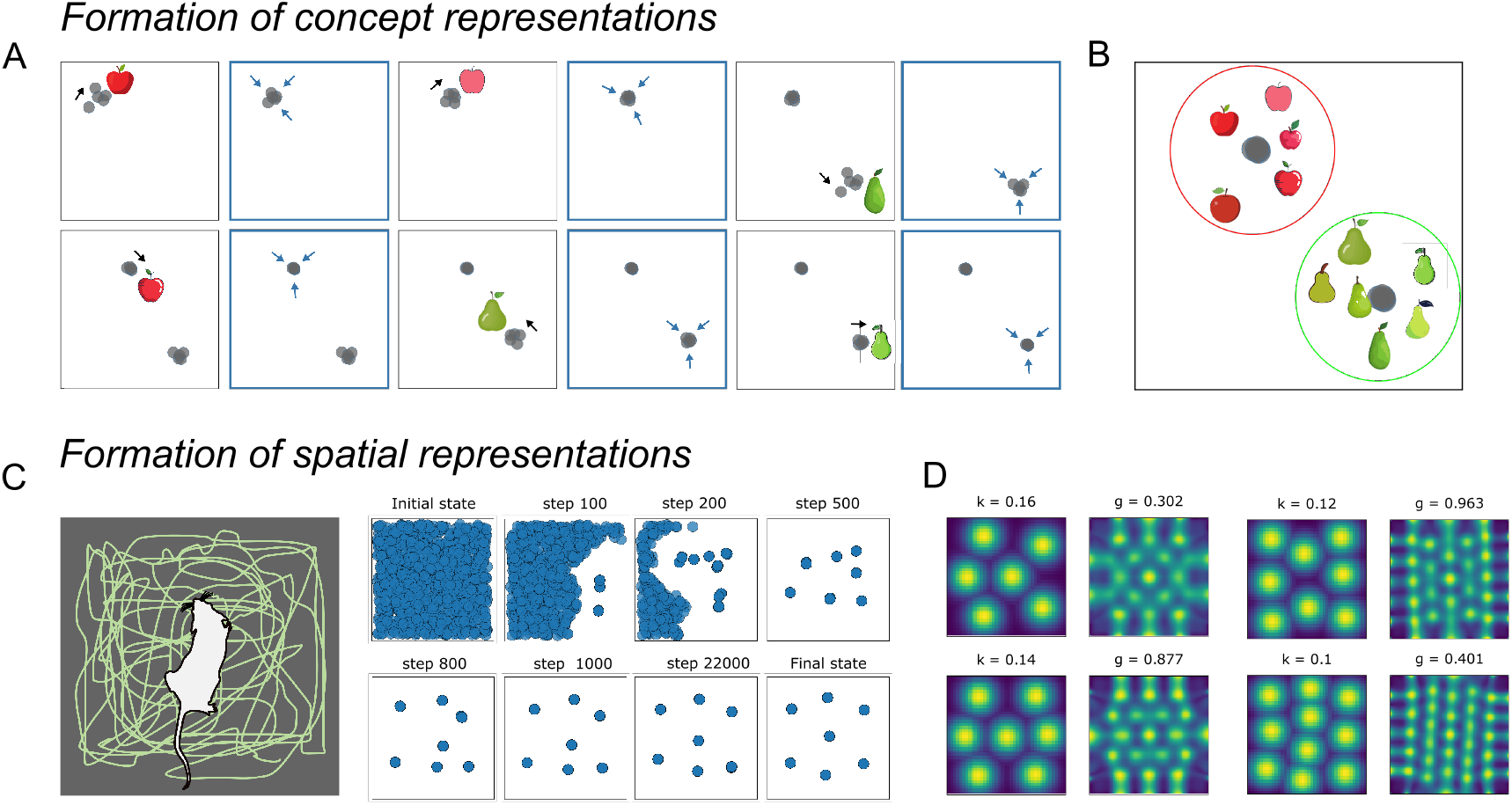
Formation of concept and spatial representations by neural flocking. A) The model learns distinct representations for Apples and Pears. The *k* winners (i.e., most activated units) adjust their receptive fields toward the current stimulus, followed by a recurrent update toward their centroid. B) This second update is sufficient to solve the coordination problem and allow SUSTAIN-d to form neural flocks or virtual clusters, which in this example represent the concepts Apple and Pear. C) Spatial representation formation. Left: The agent (e.g., a rodent) forages in an environment. Right: Development of spatial representations. Initially, SUSTAIN-d’s neuron-like units are uniformly tuned to locations. At each time step, the *k* winners both move toward the stimulus (e.g., sensory information at the current location) and each other (i.e., neural flocking). This learning dynamic creates flocks or virtual clusters of units with similar spatial tuning, akin to place-cell assemblies. These flocks tile the environment. D) Examples of grid cell-like activity patterns and corresponding spatial autocorrelograms after learning. See Fig. S1A for more examples and S1B for distributions of grid scores).

## Neural population-based model retains high-level cognitive model properties and captures concept learning behavior

One major challenge for our multi-level proposal is to account for complex behaviours that hitherto were the sole province of cognitive models. Can SUSTAIN-d with its neuron-like units account for the same behaviors that SUSTAIN does by relying on cognitive constructs? We evaluate whether SUSTAIN-d can account for human learning performance on Shepard et al.’s six classic learning problems (Fig. 3). To provide a true multi-level theory, we aim for SUSTAIN-d’s solution in terms of virtual clusters arising from neural flocking to parallel SUSTAIN’s clustering solutions, which provide a good correspondence to hippocampal activity patterns (*21*).

**Fig. 3.**
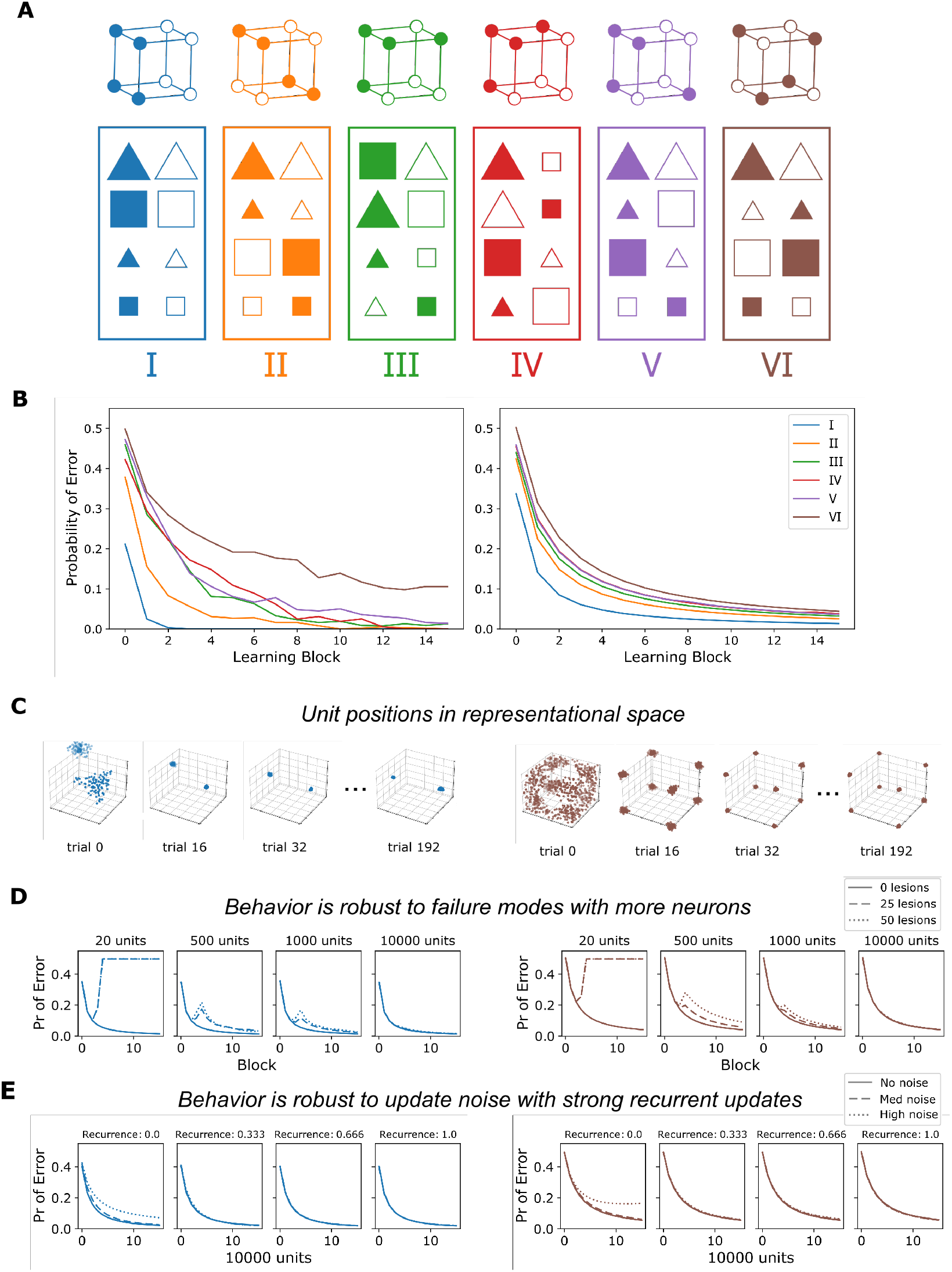
SUSTAIN-d’s brain-scale population of neuron-like units collectively displays the same behavior as the high-level cognitive model that it decomposes, while making additional predictions about robustness in neural computation. A) Six concept learning structures (*30*). Bottom: In each box, stimuli in the left and right columns are in different categories. Top: Cubes represent each stimulus in binary stimulus feature space (color, shape, size) for each structure. B) Learning curves from human behaviour (*31*) (left) and model fits (right). Probability of error is plotted as a function of learning block for each structure. C) Neuron-like units form neural flocks or virtual clusters (e.g., type I in blue and type VI in brown; see Fig. S2 for all types) that parallel the clusters in the higher-level cognitive model. The number of units are sub-sampled from the whole population for better visualization. D) The more neuron-like units, the more robust the model is when confronted by failure modes (e.g., cell death, noise, synaptic transmission failure). E) The stronger the recurrence during learning, the better the noise tolerance. See Fig. S3 for more examples.

SUSTAIN-d was trained in a manner analogous to human participants, learning through trial-and-error in these supervised learning tasks. On each trial, SUSTAIN-d, updated its neuron-like units’ positions in representational space, attention weights, and connections weights from its neuron-like to category responses (see Methods for details). All the learning updates were error driven and local, as opposed to propagating errors overs multiple network layers as in deep learning models. Whereas SUSTAIN forms a new cluster in response to a surprising error (e.g., learning that a bat is not a bird), SUSTAIN-d recruits the *k* nearest unconnected neuron-like units to the current stimulus, which is in accord with the intuition that the brain repurposes existing resources during learning. The neural flocking learning rule leads to these recruited units forming a virtual cluster.

SUSTAIN-d captured the difficulty ordering of the human learning curves (Fig. 3B, right) and its solutions paralleled those of SUSTAIN in terms of the modal number of clusters recruited (2, 4, 6, 6, 6, and 8 flocks for each of the six learning problems) and attentional allocation to features. Notably, SUSTAIN-d’s results scale to a large number of neuron-like units, producing the same output and learning curves from few (e.g. 50) to many neurons (3.2 x 10^6^ hippocampal principal cells (*28, 29*) as used here). Thus, SUSTAIN-d provides a multi-level account (*1*) of hippocampally-mediated learning that ranges from behavior to neuron-like units. SUSTAIN-d is able to display similar aggregate behavior over a wide-range of neuron-like units because its learning updates and operations can be scaled to reflect the number of units involved (see Methods).

Like the human brain, SUSTAIN-d is resistant to minor insults and faults. Each neural assembly or flock can consist of many neuron-like units (Fig. 3C, Fig. S2), not all of which are needed for the remaining units to function as a virtual cluster. SUSTAIN can be viewed as SUSTAIN-d when the number of highly activated units in response to the stimulus is 1 (i.e., *k* = 1). As *k* or total number of units increase, lesioning a subset of SUSTAIN-d’s units has negligible effects on aggregate behavior (Fig. 3D, Fig. S3A). This robustness through redundancy is consistent with operation of the brain where multiple neurons and synapses with similar properties offset the effects of damage and natural turnover of dendritic spines (*26, 32–34*).

Having multiple units combined with SUSTAIN-d’s recurrent update can also counteract noise (Fig. 3E). In these simulations, we added noise to SUSTAIN-d’s neuron-like units, which will lead to units from other assemblies (or neural flocks) becoming highly active, which can lead to incorrect decisions and disrupt learning. SUSTAIN’s recurrent update (Fig. 1C-D) ameliorates these effects of noise by pulling individual units’ responses towards the mean of flock (Fig. 3E, Fig. S3B). The same self-organizing learning rules that enables SUSTAIN-d’s neuron-like units to behave in aggregate like a cognitive model also make it more robust to noise and damage.

## Further decomposing to capture differential function in anterior and posterior hippocampus

In a multi-level mechanistic account, model components can be further decomposed to capture findings that require more fine-grain mechanisms. SUSTAIN-d decomposed SUSTAIN’s clusters into neuron-like units. Here, we further decompose SUSTAIN-d’s neuron-like units into two pools to capture functional differences between anterior and posterior hippocampus.

Anterior place fields tend to be broader and lower granularity than posterior hippocampal fields (*35*), and this appears to be a general principle at the population level (*36, 37*). For category learning studies, one prediction is that learning problems with a simpler structure that promotes broad generalization would be more anterior whereas learning problems that have a complex irregular structure would be better suited to posterior hippocampus. Indeed, this pattern holds across studies (*20, 21*) (Fig. 4A). Here, we simulate Shepard’s six learning problems, which order from what should be most anterior (Type I) to most posterior (Type VI). Type I can be solved by focusing and generalizing based on one stimulus feature, whereas Type VI requires memorizing the details of each item.

**Fig. 4.**
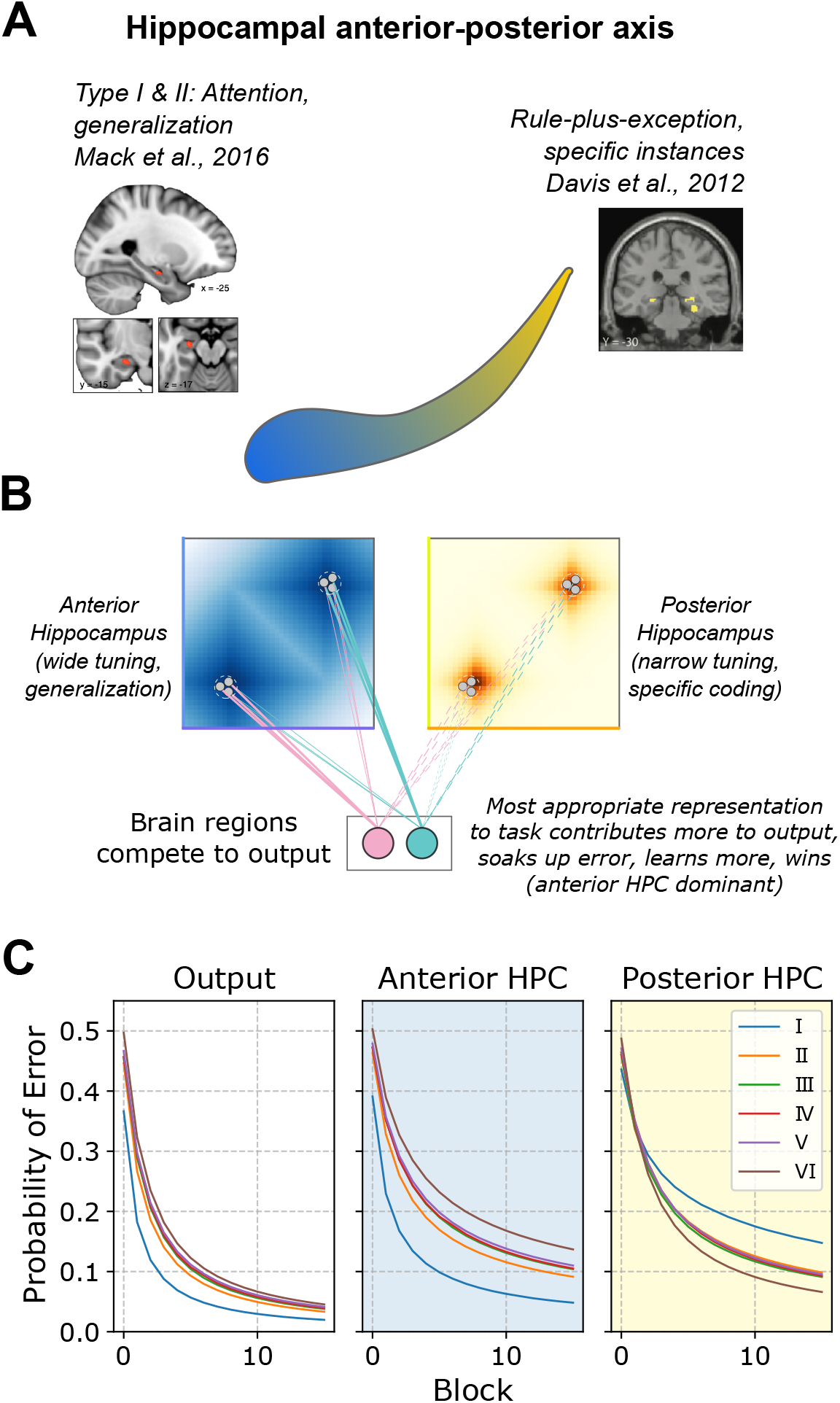
Further decomposing SUSTAIN-d to capture known functional differences along the anterior-posterior axis of the hippocampus. A) Illustration of the anterior-posterior (blue-yellow) gradient in human hippocampus. Place fields in the anterior hippocampus are broader. Likewise, anterior hippocampus is strongly activated by concept learning structures that follow broad, general rules (left), whereas posterior hippocampus is more strongly engaged by irregular rule-plus-exception structures where specific instances are important. B) SUSTAIN-d is further decomposed into a bank of units with broader tuning to model anterior hippocampus (blue) and a narrowly-tuned bank of units to model posterior hippocampus (yellow). Both banks contribute to the output and compete to exert control over the category decision. C) Model output (left) is a combination of the anterior (middle) and posterior (right) neuron-like units. The anterior units dominate for simple category structures, whereas the posterior units dominate for irregular structures.

Although the anterior-posterior distinction may best be viewed as a continuous axis, we simplified to create two banks of neuron-like units for SUSTAIN-d, one corresponding to anterior hippocampus and one to posterior hippocampus. These two banks of neuron-like units only differed in how broad their tuning was with anterior fields being broader than posterior fields. The responses from both pools of neuron-like units were pooled to determine the category response (Fig. 4B). The two pools of units formed neural flocks independently and were only linked in that they both aimed to correctly predict the category label. Thus, the pool that was more suited to a learning problem will take over by developing larger connection weights to the output units indicating the response. In effect, both pools of units compete to guide the overall response (see Methods). Thus, the anterior pool of units should guide the overall system response when a problem is very simple (e.g., Type I) whereas the posterior pool should take over when a problem is highly irregular and complex (e.g., Type VI).

As predicted, the anterior pool guided the overall response for simple problems like the Type I problem whereas the posterior pool did so for complex problems, like the Type VI problem (Fig. 4C). Indeed, the posterior pool learns the Type VI problem faster than the Type I problem because SUSTAIN-d forms eight virtual cluster (neural flocks) for the Type VI problem which is ideally suited to the narrow fields in the posterior pool which favor memorization over broad generalization, exactly the opposite functional properties of the anterior pool. These simulations suggest that the anterior-posterior axis in the hippocampus may provide complementary learning mechanisms.

## Discussion

Neuroscience is inherently a multilevel enterprise that requires multilevel theories to move from a patchwork understanding of the mind and brain toward more integrative and encompassing theories. One solution, which we advance, is to adopt a levels of mechanism approach (cf. Craver) in which the components of a higher-level model are unpacked into their own mechanisms. Here, we offered a multilevel explanation of category learning in which a cognitive model at the top level that accounts for behavior is decomposed into populations of neuron-like units that form assemblies.

This breakthrough was largely achieved by a novel learning rule that promoted neural flocking in which neuron-like units with related receptive field properties became more similar by coordinating using recurrence. Like a flock of birds, these units functioned as a collective, providing an account of how symbols and other cognitive constructs, such as clusters, can arise from neuron-like computing elements following biologically consistent operations.

The recurrent update in the learning rule was inspired by biological recurrence in the hippocampal formation (within the hippocampus and big-loop recurrence in the medial temporal lobe; (*25*)), where multiple passes allow for deeper processing. Symbol-like neural representations naturally form through our implementation of recurrence, suggesting that functional neural assemblies can form through a flocking mechanism like in birds. With recent advances in large-scale neural recordings over time, future neurophysiological or imaging studies in animals that record from hippocampus across time and over learning could search for such a neural mechanism.

With a neural population and a recurrent mechanism, the model also naturally captures the brain’s tendency to encode the same information in many neurons (i.e., redundancy; (*32–34*)), which makes the system more tolerant to neuronal damage, natural synaptic turnover, or noise. Notably, there could be more than one recurrent update, as there are multiple recurrent loops in the brain. Future work could introduce more recurrent steps or different forms of recurrence such as constraining it by known anatomical pathways.

It is worth noting that the model assumes a sparse representation where most neurons are dormant but active neurons are highly active for a small group of stimuli, which parallels sparse coding in the hippocampus. Furthermore, the sparse code of a fixed proportion of highly-activated winners in a neural population is consistent with recent hippocampal neurophysiological evidence showing that the overall activity of place cells within an environment is constant (*38*). This work shows how localist coding models (i.e., with sparse, highly-tuned neurons) can be implemented in a neural population (*39–42*). One way for related approaches to show mixed selectivity is for different stimulus dimensions to be relevant in different contexts. For example, a neuron may respond to wool socks in a clothing context and to tart flavours in a food context with the relevant dimensions or features in each context being non-overlapping.

One benefit of multi-level theories is that model components can be decomposed into their own mechanisms as desired. In effect, one can selectively zoom in to consider the aspects of the mechanism of interest at a finer grain. In the last set of simulations, we further decomposed SUSTAIN-d’s neuron-like units into two banks of units corresponding to anterior and posterior hippocampus. Further decomposing SUSTAIN-d allows us to account for finer-grain phenomena and make new predictions.

We found that the bank of units best suited to the task dominated learning. For learning problems with a simple structure, the broader receptive fields of the anterior bank dominated. When the learning problem had an irregular structure that required memorization, the posterior bank dominated. By varying the broadness of the receptive fields, we introduced a generally applicable framework in which modules compete with one another to guide behavior but are ultimately complementary in terms of accomplishing the goals of the overall system. This cross-region competitive framework captures the common finding across cognitive neuroscience where brain regions that have the more appropriate or useful representations for the task at hand are more strongly activated and contribute more to behavior.

In the future, we plan to extend the method to include connections across modules (e.g., excitatory/inhibitory) based on known anatomy and functional properties, and to model more interacting brain regions. The neuron-like units themselves could be further decomposed and elaborated to behave more like biological neurons. For our present purposes, we did not require spiking neurons with complex dynamics. However, just as SUSTAIN was decomposed into SUSTAIN-d, so too can SUSTAIN-d be decomposed as desired, all the while retaining the overarching principles and behavior of the higher-level models.

In sum, cognitive neuroscience can benefit from multi-level explanations by exploring and bridging mechanisms across levels. We have many cognitive models that characterize behavior successfully, but are in need to be decomposed into a set of mechanistic processes that could be implemented in the brain. In recent years, neuroscience is finally putting more emphasis on behaviour (*5*), but we suggest that for a complete account of the cognitive function of interest, a successful high-level explanation of the behavior (e.g., through a cognitive model) that can be decomposed into the relevant lower-level mechanisms is key.

## Acknowledgements

Funding: This work was supported by the Medical Research Council UK (SUAG/045 G101400) and a Leverhulme Trust Early Career Fellowship (Leverhulme Trust, Isaac Newton Trust: SUAI/053 G100773, SUAI/056 G105620, ECF-2019-110) to R.M.M, and the Wellcome Trust (WT106931MA) and a Royal Society Wolfson Fellowship (18302) to B.C.L. Authors contributions: R.M.M.: conceptualization, data curation, formal analysis, funding acquisition, methodology, software, visualization, writing – original draft, writing – review & editing. B.C.L.: conceptualization, formal analysis, funding acquisition, methodology, resources, supervision, visualization, writing – original draft, writing – review & editing. Competing interests: None. Data and materials availability: Code and simulation results will be available at https://github.com/robmok/multiunit-cluster and at OSF at time of publication. Fruit images were obtained from vecteezy.com. For the purpose of open access, the author has applied a Creative Commons Attribution (CC BY) licence to any Author Accepted Manuscript version arising from this submission.

## Methods

### Overview and motivation of the model

As discussed in the main text, SUSTAIN-d is a decomposition of SUSTAIN (*18*), a cognitive model of concept learning that has captured behaviour (*18, 19*) in a number of tasks and brain-activity patterns including in the hippocampus and medial temporal lobe structures (*16, 20, 21*) (also see (*43*)). The formal specification of SUSTAIN is included in the aforementioned papers. Whereas SUSTAIN contains clusters (a cognitive construct) that are recruited in response to surprising events, SUSTAIN-d decomposes the notion of cluster into neuron-like units that coordinate to form virtual clusters through a flocking learning rule as described below.

### Model Learning Implementation details

The model was initialized with a population of neuron-like units. Full-scale simulations used 32,000,000 units to model the hippocampus principal cell population. To determine the best fitting parameters for these simulations (see below), 10,000 units were used to reduce computational costs. Units were placed randomly (uniformly) in the stimulus feature space, where all units are inactive or unconnected to the task context. On the first trial (no output), or when the model makes an error (greater output for the incorrect category), *k* (proportion of total; set to 0.00005 or 0.005% for the hippocampus simulation and 0.01 for parameter search) neurons are recruited at the current stimulus’ position (note that number of units and *k* does not change model behavior, see section on scaling below). Once units are recruited, they are connected and activate in response to stimulus input, and their activations contribute to the category decision. On each trial, the winners’ activations contribute to the output decision, and they update their tuning by moving toward the current stimulus (Kohonen learning rule) and then towards their own centroid (recurrent update), and the attention weights and connection (i.e., output) weights are updated through learning.

Specifically, the model takes a stimulus vector as input on each trial, and the most strongly activated *k* connected neuron-like units are considered winners (k-winners-take-all; k-WTA) and the activation of these units are computed on each trial:

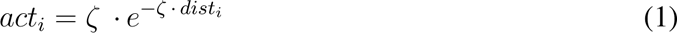

where *ζ* is a positive scaling parameter which controls the steepness of the tuning curve, and *dist_i_* is the attention-weighted distance between neuron *i*’s position *pos_i_* and the stimulus *x* in the *R^n^* representational space they inhabit:

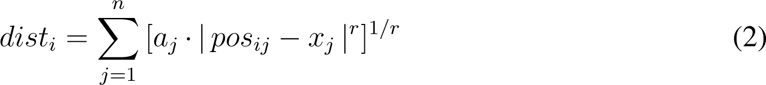

where *r* is set to 1 for the city-block distance (for separable-dimension stimuli) and the non-negative *a_j_*attention weights sum to 1. The *n* attention weights correspond to each stimulus feature dimension, and their magnitude reflects the importance of each feature. The attention weighting also corresponds to each unit’s receptive field, which is centered on its position along each feature dimension (Fig. 1B, attention weighting). *ζ* controls the steepness of the receptive field and will be used to model the different tuning properties of the anterior and posterior hippocampus (see below). As such, units most strongly tuned to the current stimulus input are activated and the activity is propagated forward to produce the categorization decision. Only winners have non-zero output that contribute to the category decision:

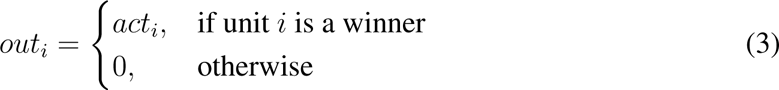

and evidence for category decisions propagates from the output:

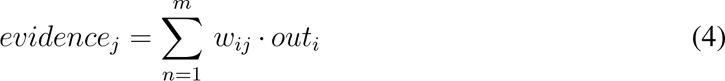

where *w_ij_* is the connection weight (Fig. 1B, cyan and pink connections) between unit *i* and decision *j* and *m* is the number of units. Finally, the probability of making decision *j* is computed by:

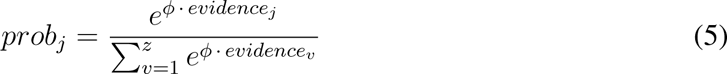

where *z* is the number of possible decisions and *φ* is a non-negative decision certainty parameter, or inverse temperature (see (*18, 44*) for related formulations). If there are no connected units (e.g. first trial) or fewer than *k* winners, then no units respond or fewer than *k* units respond, respectively.

During learning, the *k* winners update twice, once toward the stimulus and a second time toward each other, which supports neural flocking. We view unit updating as a continuous process through time relying on recurrent connections, which we simplify here to two simple updates. In the first update, the *k* winners update their positions toward the current stimulus’ position on each trial according to the Kohonen learning rule:

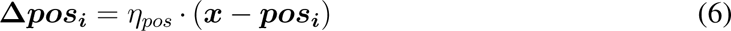

where *η_pos_* is the learning rate, *x* is the current stimulus’ location and *pos_i_* is to unit *i*’s position in representational space (Fig. 1C, 1st update). Bold type is reserved for vectors. This is the first learning step, where the initial forward pass of stimulus information occurs and the first clustering update is applied. In the second update, the *k* winners perform an additional recurrent step which re-codes the stimulus based on their activity, updating their positions toward the centroid of the *k* winners’ positions:

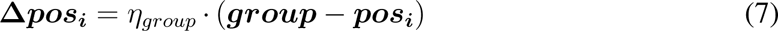

where *η_group_* is the learning rate for the recurrent update and ***group*** is the centroid (mean position) of all the winners (Fig. 1C, 2nd update). With this double-update rule, neurons update their tuning profiles to activate more to stimuli that inhabit that part of space in a coordinated fashion, and over the course of learning will typically stabilize into a portion of the space adaptive for the task. As co-activated units cluster together and become more similar to each other in tuning, this group naturally lead to a virtual cluster (i.e., many units that are similar tuned to a concept) similar to a cluster (*18*) and akin to to a hippocampal place or concept cell assembly where assemblies of neurons show similar tuning and co-activate to similar stimuli or environmental features (*7–9*).

To update attention weights over learning, the model applies gradient ascent on the summed activation of winning units minus the summed activation of the non-winner units (i.e., connected but not a winner) with learning rate *η_attn_*. Note that this learning rule is local (i.e., no backpropagation).

Connection weights are updated using descent on error (based on the output *prob_j_*) using cross-entropy loss with one-hot target vector (i.e., one category is correct) on each trial, with learning rate *η_cweights_*. To display the output of the model and for fitting to the behavioural data, we plot 1 *− prob_j_* (where *j* is the correct category decision), averaged for each block to match the error learning curves as done in prior work.

As the model activations, outputs, and learning update values will vary depending on the number of units, we scaled the learning rates to retain consistent outputs across models with different number of units. This scaling means that changing the total number of units and *k* (winners; proportion of total) does not change the learning behavior and output of the model, meaning the model can scale from a small model with a few neurons to millions of neurons without losing its theoretical essence and cognitive capacities, allowing us to bridge across levels (assuming no noise – see main text for beneficial effects of a larger number of neuron-like units with noise). Furthermore, this allowed us to perform parameter search for fitting human behavior with fewer units and different *k*, whilst reporting the hippocampal-scale simulation with many more units. First, the learning rate of the connection weights was divided by *k*, so that weight updates would scale by the *k* winners that contribute to the output on each trial. As the attention weights are updated locally by gradient ascent to the winner neurons’ activations relative to the loser neurons’ activations, we divided the update (the gradient) by the number of active units (i.e. total number of the winner and loser neurons included to compute the gradient).

### Model fitting: human concept learning behavior

To model the classic Shepard, Hovland, & Jenkins (*30*) results, we fit the learning curve data (minimizing sum of squared errors) from the Nosofsky et al. (*31*) replication of the Shepard et al. study. We present a brief overview of this classic study here. Participants learned to categorize eight stimuli that varied on three binary feature dimensions (shape, size, and color) into two categories. The concept structure was one of six possible logical structures from Shepard et al. (Fig. 3A). On each trial, participants categorized each stimulus into a category and was provided feedback, learning by trial and error. Participants completed blocks of 16 trials (with two repetitions of each stimulus). Participants continued learning until they made no errors in four sub-blocks of eight trials, or if they completed 25 blocks (400 trials). In both studies, they plotted error curves (1 *−* proportion correct) for the first 16 blocks (16 trials per block), which are the data we will fit (Fig. 3B, left).

Task blocks consisted of 16 stimuli presented in a randomized order. To obtain error curves for each parameter set, a random stimulus sequence for each problem type was generated 25 times, and the error curves were produced by taking the mean across those iterations. To maintain consistency, each iteration was seeded with a specific number, so that the 25 sequences were the same across the different parameters.

Model learning curves were fit to display the human pattern of results. We performed a hierarchical grid search across the parameters. For the standard model (i.e. no separation of brain regions or modules), there were 6 free parameters: *ζ*, *φ*, and four learning rates (attention weights *η_attn_*, connection weights *η_cweights_*, Kohonen update *η_pos_*, recurrent update *η_group_*). To fit the model that includes a separate anterior and posterior bank of units, separate tuning (*ζ*) parameters were used for each bank with the constraint that the anterior bank should have a broader tuning than the posterior bank of units (12 free parameters).

### Robustness to failure modes: noise and lesion experiments

To demonstrate the beneficial effect of having a population of neurons (rather than a single unit or cluster in cognitive models) and the recurrent update, we simulated Shepard’s problems with different failure modes during the learning process and how robust the model was to these perturbations.

To simulate noise in the learning process, we added noise to the units’ positions. For each trial, noise was sampled from a *n* dimensional Gaussian distribution (corresponding to *n* features) with zero mean and standard deviation of 0, 0.5, or 1.0 which was added to the update. By adding noise to the unit’s position in representational space, this causes potential problems for 1) selecting the appropriate *k* neurons as winners, 2) appropriate updating of the attention weights, and 3) appropriate updating of the connection weights.

To simulate damage-like events in the neural population, we performed a lesion-like experiment where we randomly removed a subset of the active neurons from the model, simulating typical biological changes such as neuron death or synaptic turnover. For a simple illustration of the beneficial effect of the number of neurons on damage-like events, we set up one “lesion” event at trial 60 where 0, 25, or 50 units were removed and rendered inactive from that point on. The results hold with more lesion events or a larger number of neurons removed.

### Unsupervised learning on spatial tasks

To simulate a rodent foraging in an environment, we placed an agent in a two-dimensional square environment and produced a randomly-generated 500,000 set of steps with the restriction that the agent could not step out of the environment. On each trial, it was able to move left, right, up, or down in steps of 0, 0.025, 0.05, or 0.075. The environment was a square that spanned from 0 to 1 on the horizontal and vertical dimensions.

For unsupervised learning, the model could recruit units like SUSTAIN does by relying on a surprise signal. Here, we further simplify as in (*16*) and assume all units in the population are relevant to the current context. Unit positions were updated according to the learning rules specified above. While a rodent’s actual environment contains many features, we assumed these features effectively reduce to a two-dimensional space corresponding to coordinates within the agent’s enclosure.

On each trial, the agent moved a step (randomly selected over four directions and four step sizes; one trial), and the model updated the *k* winners with the Kohonen learning rule as before, with an annealed learning rate so that the units would eventually settle and stabilize into a particular location:

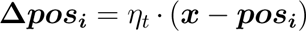

where *η_t_* is the learning rate at time *t*. The learning rate followed an annealing schedule:

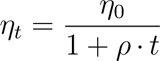

where *η*_0_ is the initial learning rate and *ρ* is the annealing rate set to 4 *×* 10*^−^*^12^ (see (*16*)). The recurrence update learning rate *η_group_* was fixed at 1.0, though smaller values such as 0.8 and 0.6 produce similar results.

To compute grid scores at the end of learning, activation maps were produced by generating a new movement trajectory with 250,000 steps (as above), and computing the unit activations based on their positions at the end of learning (i.e. freezing the positions; see (*16*)). For each value of *k*, we ran 100 simulations and computed the grid scores at the end of learning. The activation maps were binned in 40×40 bins (original 100×100), then normalized by the number of visits to each binned location (normalized activation map). Grid scores were calculated based on (*45*). Briefly, the spatial autocorrelogram of the activation maps were calculated as defined in (*46*), and gridness was computed using the expanding gridness method, where a cicular annulus with a radius of eight bins was placed on the center of the autocorrelogram, with the central peak removed. The annulus was rotated in 30° steps, and the Pearson correlation between the rotated and unrotated version of the spatial autocorrelogram was recorded. The highest correlation value for 30°, 90°, and 150° rotations was subtracted from the lowest correlation value at 0°, 60°, and 120° to give an interim grid score. This was repeated expanding the annuls by two bins, up to 20. The final grid score was the highest interim grid score.

## Supplementary Figures

**Fig. S1.**
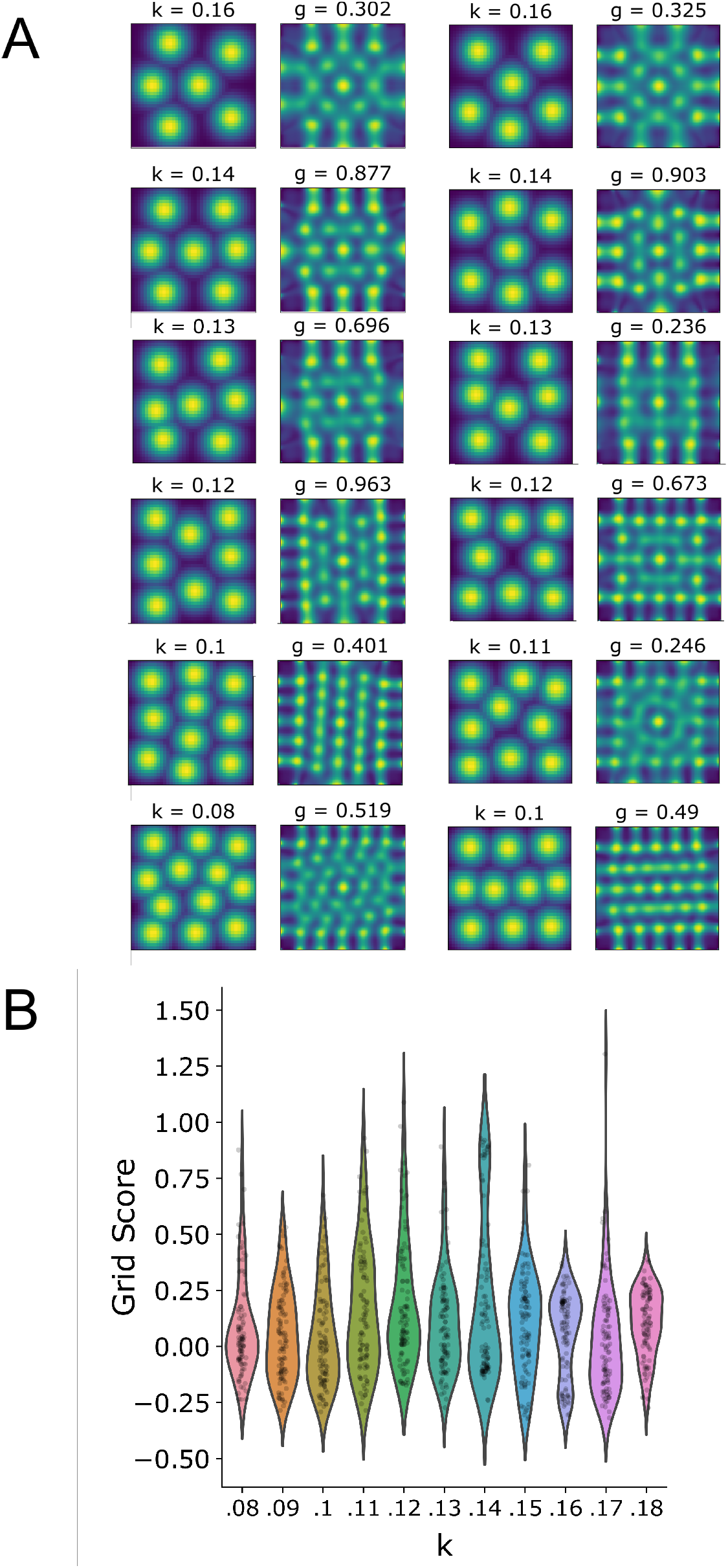
SUSTAIN-d develops grid-cell like activity patterns with learning. A) Examples of grid cell-like activity patterns and corresponding spatial autocorrelograms after learning across different values of *k*. B) Distribution of grid scores across different values of *k*. The recurrent strength is set to 1.0 but similar results are obtained with values of 0.8 and 0.6. *k* controls the number of flocks or spatial cell assemblies that form.

**Fig. S2.**
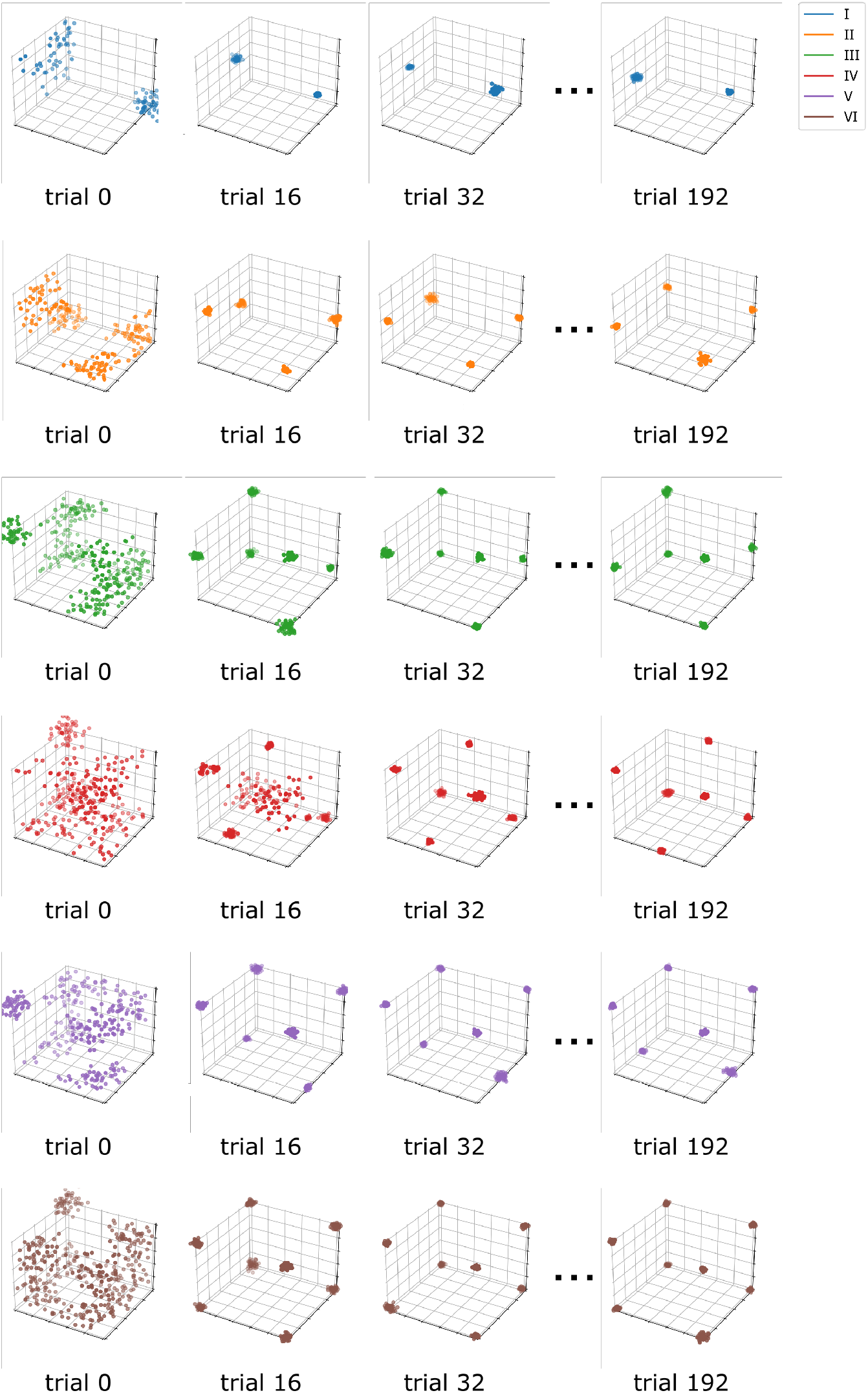
Positions of neuron-like units over learning for each concept learning structures from Type I to Type VI (top to bottom) in SUSTAIN-d. To improve visualization, the number of units are sub-sampled from the full population. As noted in the main text, neural flocks or virtual clusters form that parallel the number and form of clusters in the higher-level cognitive model SUSTAIN.

**Fig. S3.**
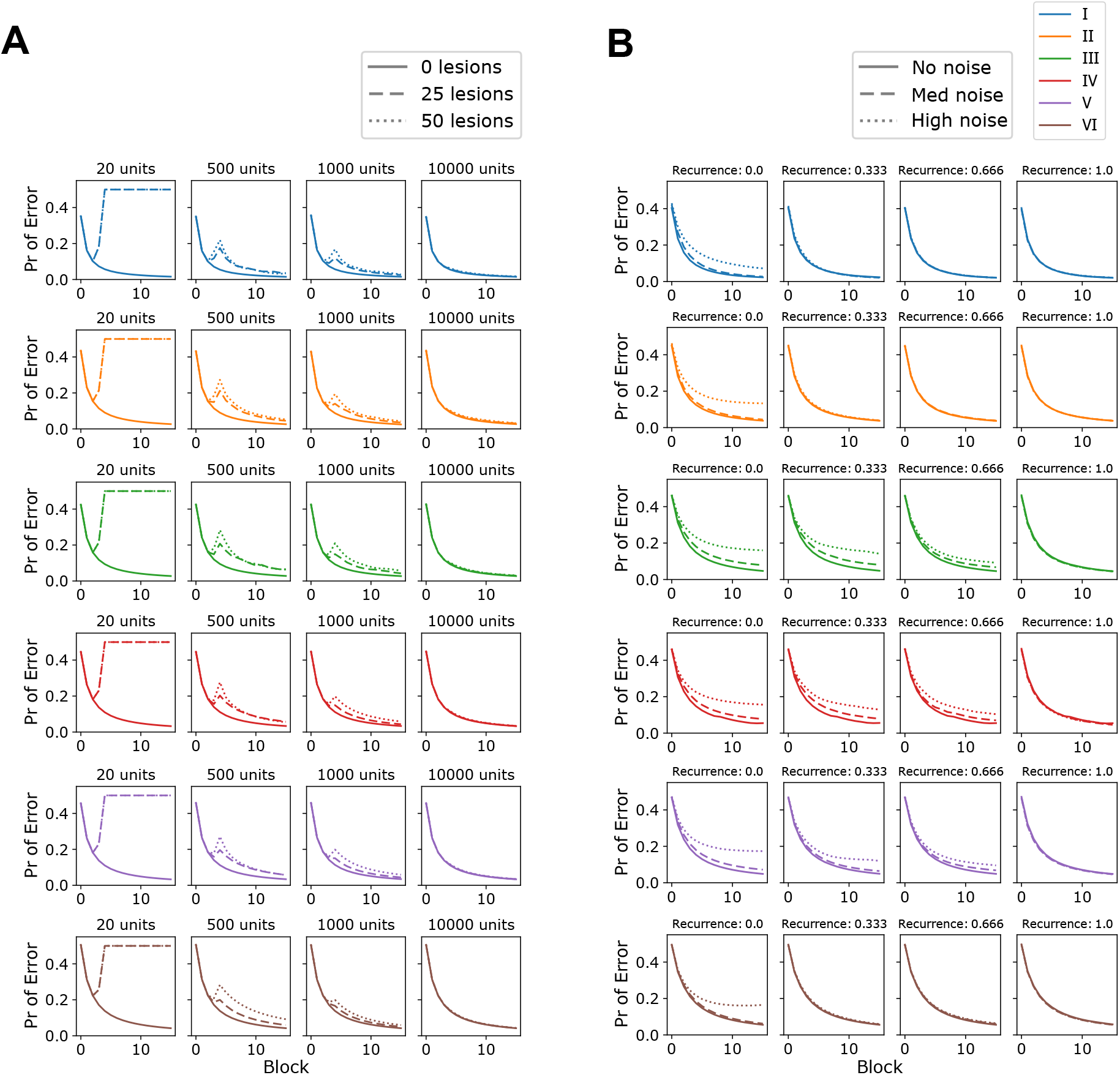
SUSTAIN-d predicts that having many neuron-like units forming neural assemblies (i.e., neural flocks or virtual clusters) makes the model robust to individual units failing and noise across all units. With a large number of units, SUSTAIN-d’s aggregate behavior is not noticeably affected by these challenges. A) Lesion simulations. In these simulations, a ‘lesion’ event occur at trial 60 where 0, 25, or 50 units were dropped out of the model. For each concept learning problem, the learning curves are plotted for each model size (i.e., number of units in the model) simulation. The more total units there are providing redundancy and stability, the more robust the model is to lesions. This result holds across all concept learning problems for both measures. B) Noise simulations. Recurrence makes SUSTAIN-d more robust to noise. For each concept learning problem, the learning curves are plotted for each recurrent strength. In this simulation, Gaussian noise is added to the position of the units on each trial after the first update step (low=0, med=0.5, high=1.0 s.d.). A recurrent step (second update in the learning rule), helps reduce the effect of noise by pulling individual units’ responses towards the mean of flock or the virtual cluster. The greater the recurrent strength, the more the effect of noise is suppressed across all concept learning problems. Each simulation used 10,000 units but the results generalize to different number of units.

## Notes

### Competing Interest Statement

The authors have declared no competing interest.

### Summary of Updates

Reverting back to original preprint (as journal doesn't allow updating after peer review)

## References

1. C. F. Craver, Explaining the Brain (Oxford University Press, 2007).

2. D. Marr, T. Poggio, AI Memo 357 (1976).

3. B. C. Love, Philosophical Transactions of the Royal Society B: Biological Sciences 376, 20190632 (2021).

4. J. Bickle, Topics in Cognitive Science 7, 299 (2015).

5. J. W. Krakauer, A. A. Ghazanfar, A. Gomez-Marin, M. A. MacIver, D. Poeppel, Neuron 93, 480 (2017).

6. D. Hebb, The Organization of Behavior: A Neuropsychological Theory (Wiley, New York, 1949).

7. G. Buzsáki, Neuron 68, 362 (2010).

8. H. Eichenbaum, Neuroscience Letters 680, 88 (2018).

9. R. Quian Quiroga, L. Reddy, G. Kreiman, C. Koch, I. Fried, Nature 435, 1102 (2005).

10. K. D. Harris, J. Csicsvari, H. Hirase, G. Dragoi, G. Buzsáki, Nature 424, 552 (2003).

11. J. M. O’Keefe, L. Nadel, J. O’Keefe, The hippocampus as a cognitive map (Clarendon Press, Oxford, 1978).

12. E. I. Moser, E. Kropff, M.-B. Moser, Annual Review of Neuroscience 31, 69 (2008).

13. A. Horner, J. Bisby, E. Zotow, D. Bush, N. Burgess, Current Biology 26, 842 (2016).

14. J. L. S. Bellmund, P. Gärdenfors, E. I. Moser, C. F. Doeller, Science 362, eaat6766 (2018).

15. T. E. Behrens, et al., Neuron 100, 490 (2018).

16. R. M. Mok, B. C. Love, Nature Communications 10, 5685 (2019).

17. E. C. Tolman, Psychological Review 55, 189 (1948).

18. B. C. Love, D. L. Medin, T. M. Gureckis, Psychological Review 111, 309 (2004).

19. B. C. Love, T. M. Gureckis, Cognitive, Affective, & Behavioral Neuroscience 7, 90 (2007).

20. T. Davis, B. C. Love, A. R. Preston, Cerebral Cortex 22, 260 (2012).

21. M. L. Mack, B. C. Love, A. R. Preston, Proceedings of the National Academy of Sciences 113, 13203 (2016).

22. M. L. Mack, A. R. Preston, B. C. Love, Nature Communications 11, 46 (2020).

23. M. B. Broschard, J. Kim, B. C. Love, E. A. Wasserman, J. H. Freeman, Neurobiology of Learning and Memory 185, 107524 (2021).

24. C. W. Reynolds, Proceedings of the 14th annual conference on Computer graphics and interactive techniques - SIGGRAPH ‘87 (ACM Press, Not Known, 1987), pp. 25–34.

25. R. Koster, et al., Neuron 99, 1342 (2018).

26. A. Attardo, J. E. Fitzgerald, M. J. Schnitzer, Nature 523, 592 (2015).

27. M. L. Mack, B. C. Love, A. R. Preston, Neuroscience Letters 680, 31 (2018).

28. G. S^̌^imić, I. Kostović, B. Winblad, N. Bogdanović, The Journal of Comparative Neurology 379, 482 (1997).

29. N. Sukenik, et al., Proceedings of the National Academy of Sciences 118, e2018459118 (2021).

30. R. N. Shepard, C. I. Hovland, H. M. Jenkins, Psychological Monographs: General and Applied 75, 1 (1961).

31. R. M. Nosofsky, M. A. Gluck, T. J. Palmeri, S. C. Mckinley, P. Glauthier, Memory & Cognition 22, 352 (1994).

32. H. Barlow, Network: Computation in Neural Systems 12, 241 (2001).

33. N. S. Narayanan, Journal of Neuroscience 25, 4207 (2005).

34. J. L. Puchalla, E. Schneidman, R. A. Harris, M. J. Berry, Neuron 46, 493 (2005).

35. M. Jung, S. Wiener, B. McNaughton, The Journal of Neuroscience 14, 7347 (1994).

36. J. Poppenk, H. R. Evensmoen, M. Moscovitch, L. Nadel, Trends in Cognitive Sciences 17, 230 (2013).

37. I. K. Brunec, et al., Current Biology 28, 2129 (2018).

38. S. Tanni, W. de Cothi, C. Barry, bioRxiv (2021).

39. J. S. Bowers, Psychological Review 116, 220 (2009).

40. D. C. Plaut, J. L. McClelland, Psychological Review 117, 284 (2010).

41. R. Quian Quiroga, G. Kreiman, Psychological Review 117, 291 (2010).

42. J. Sučević, A. C. Schapiro, A neural network model of hippocampal contributions to category learning, *preprint*, Neuroscience (2022).

43. C. R. Bowman, T. Iwashita, D. Zeithamova, eLife 9, e59360 (2020).

44. J. K. Kruschke, Psychological Review 99, 22 (1992).

45. A. Banino, et al., Nature 557, 429 (2018).

46. F. Sargolini, et al., Science 312, 758 (2006).

